# Quantifying biosynthetic network robustness across the human oral microbiome

**DOI:** 10.1101/392621

**Authors:** David B. Bernstein, Floyd E. Dewhirst, Daniel Segrè

## Abstract

Metabolic interactions, such as cross-feeding, play a prominent role in microbial community structure. For example, they may underlie the ubiquity of uncultivated microorganisms. We investigated this phenomenon in the human oral microbiome, by analyzing microbial metabolic networks derived from sequenced genomes. Specifically, we devised a probabilistic biosynthetic network robustness metric that describes the chance that an organism could produce a given metabolite, and used it to assemble a comprehensive atlas of biosynthetic capabilities for 88 metabolites across 456 human oral microbiome strains. A cluster of organisms characterized by reduced biosynthetic capabilities stood out within this atlas. This cluster included several uncultivated taxa and three recently co-cultured *Saccharibacteria* (TM7) phylum species. Comparison across strains also allowed us to systematically identify specific putative metabolic interdependences between organisms. Our method, which provides a new way of converting annotated genomes into metabolic predictions, is easily extendible to other microbial communities and metabolic products.

## Introduction

Metabolism, in addition to enabling growth and homeostasis for individual microbes, is a powerful “currency”, that contributes to the organization of microbes into complex, dynamic societies. Metabolic interactions are believed to influence microbial community structure and dynamics at multiple spatial and temporal scales^1-5^. For example, through cross-feeding, a compound produced by one species might benefit another, leading to a network of metabolic interdependences^5-10^. An extreme case of interdependence between microbes is believed to underlie what is usually known as “microbial uncultivability”^11^, i.e. the fact that many microbes isolated from a given environment do not grow in pure culture on standard laboratory conditions. This observation, originally proposed as “the great plate count anomaly”^12^, has motivated interest in understanding the possible mechanisms underlying unculturability^11,13,14^. One class of mechanisms is based on the concept that the growth of uncultivable microbes depends on their community context via diffusible metabolites produced by their neighbors^14^. These dependent microbes are often referred to as fastidious, due to their limited biosynthetic capabilities and reliance on externally supplied metabolites for growth. The prominence of fastidious microbial organisms across the tree of life and their potential importance in microbial community structure is highlighted by the recent identification of the candidate phyla radiation – a large branch of the tree of life consisting mainly of uncultivated organisms with small genomes and unique metabolic properties^15-17^.

Some of the most promising strides in understanding metabolic interdependences between microbes have been taken in the study of the human oral microbiome. The human oral microbiome serves as an excellent model system for microbial communities research, due to its importance for human health and ease of access for researchers^18,19^. For example, the order of colonization of species and the spatial arrangement of microbes in dental plaque have been thoroughly characterized^20,21^. The human oral microbiome consists of roughly 700 different cataloged microbial species, identified by 16S rRNA microbiome sequencing^18,22^. Importantly, 63% of species in the human oral microbiome have been sequenced, including several uncultivated and recently-cultivated strains that have implications in oral health and disease^23,24^. Exciting recent work has led to successful laboratory co-culture growth of three previously uncultivated organisms, the *Saccharibacteria* (TM7) phylum taxa: *Saccharibacteria* bacterium HMT-952 strain TM7x^25,26^, *Saccharibacteria* bacterium HMT-488 strain AC001 (not yet published), and *Saccharibacteria* bacterium HMT-955 strain PM004 (not yet published). *Saccharibacteria* are prominent in the oral cavity and relevant for periodontal disease^27,28^. Due to their importance, they were among the first uncultivated organisms from the oral microbiome to be fully sequenced via single-cell sequencing methods^29^, and represent the first cultivated members of the candidate phyla radiation^25^. Thus, their metabolic and phenotypic properties are of great interest for oral health and microbiology in general.

In parallel to achieving laboratory growth of uncultivated bacteria, a major unresolved challenge is understanding the detailed metabolic mechanisms that underlie their dependencies. Ideally, one would want to computationally predict, directly from the genome of an organism, its biosynthetic capabilities and deficiencies, so as to translate sequence information into phenotypes, mechanisms, and community-level predictions^30^. A number of approaches, based on computational analyses of metabolic networks, have contributed significant progress towards this goal^31-33^, including in the context of microbial communities^4,5,33-42^. At the heart of these methods are metabolic network reconstructions, formal encodings of the stoichiometry of all metabolic reactions in an organism, that are readily amenable to multiple types of *in silico* analyses and simulations^44^. Recent exciting progress has led to the automated generation of “draft” metabolic network reconstructions for any organism with a sequenced genome^45^, opening the door for the quantitative study of large and diverse microbial communities. Despite this promise, the most commonly used metabolic network analysis methods, such as flux balance analysis (FBA)^46^ or its dynamic version (dFBA)^47^, are not applicable to these draft metabolic networks due to gaps (missing or incorrect reactions) in the metabolic network. Methods for “gap-filling” draft reconstructions can alleviate this problem at the expense of an increased risk for false positive predictions. Additionally, gap-filling requires specific assumptions on the growth media composition – which is often difficult to obtain for diverse environmental isolates and by definition unknown for uncultivated organisms. Thus, the capacity to provide predictions based on unelaborated genome annotation, and on limited knowledge about an organism’s growth environment remains an important open challenge. Metabolic network analysis methods that are less dependent on gap-filling have been applied to the analysis of draft metabolic reconstructions, generally with a focus on metabolic network topology^48-50^. Some of these methods have provided valuable insight into the biosynthetic potentials of organisms and metabolites^51,52^, the chance of cooperation or competition between species^53-56^, and the relationship between organisms and environment^48,57,58^ including in the human gut microbiome^59^. However, these methods often depend on specific assumption on environmental conditions^49,50^, or cannot be easily reconciled with stoichiometry-based constraints^48^.

Here we introduce a new method, which alleviates the above limitations, and provides a novel metabolic prediction – an estimate of biosynthetic network robustness. Our method applies a probabilistic approach to define and compute a metric that provides an estimate of which metabolites, such as biomass components, are robustly synthesized by a given metabolic network and which would likely need to be supplied from the environment/community. Discrepancies in these calculated estimates between organisms can be used to generate hypotheses regarding microbial auxotrophy and metabolic exchange in microbial communities. Importantly, our metric can provide an environment-independent characterization by randomly sampling many different possible nutrient combinations, and is not dependent on *a priori* biosynthetic pathway definitions as it depends only on the stoichiometric constraints of the metabolic network. We applied this method to a large number of organisms from the human oral microbiome, and identified broad trends in biosynthetic capabilities. We focused in particular on uncultivated microorganisms, including three recently co-cultured *Saccharibacteria* (TM7) strains. In addition to highlighting their biosynthetic deficiencies, we developed specific hypotheses for their metabolic exchange with growth-supporting co-culture partners.

## Analysis Method

Our newly developed method quantifies a concept we call biosynthetic network robustness. Robustness, in this sense, refers to the ability of the network to produce a specified target metabolite, from variable metabolic precursors. In essence, our metric for biosynthetic network robustness provides a measure of how well a particular metabolic network can produce a particular target across a uniformly sampled set of possible environments.

The inspiration for this method comes from the statistical physics concept of percolation. Percolation theory has been applied broadly with applications ranging from materials science to epidemiology, as well as to the study of cascading metabolic failure upon gene deletions in metabolism^60^. In percolation theory the robustness of a network can be characterized by randomly adding or removing components (nodes or edges) of a network and assessing network connectivity^61^. We utilized this concept to characterize the network robustness of a particular metabolic network towards a specified target metabolite by randomly adding input metabolites to the network and assessing the network’s ability to produce the specified target metabolite.

To implement our method, we first introduced a probabilistic framework for analyzing metabolic networks (Figure 1 A). In this framework, every metabolite can be considered to be drawn from a Bernoulli distribution, *i.e.* present in the network with a given input probability (*P_in_*). These probabilities could represent beliefs about the environment, chances of metabolites being available from a host organism, or any arbitrary prior on metabolite inputs. Throughout the implementation of our method, we have assigned *P_in_* to be an identical value for all input metabolites. However, future implementations of this probabilistic framework could easily utilize *P_in_* values that vary across metabolites, e.g. matching experimentally measured abundances. Following the assignment of *P_in_*, the network structure can be used to calculate the output probability (*P_out_*) of some specified target metabolite. In practice, random sampling of probabilistically drawn input metabolite sets is used to calculate the probability of producing the target metabolite. For each random sample, flux balance analysis^46^ with inequality mass balance constraints is used to assess the networks ability to produce the target metabolite (for a complete explanation of how flux balance analysis is implemented in this context, see methods section: Algorithm functions, *feas*).

**Figure 1.**
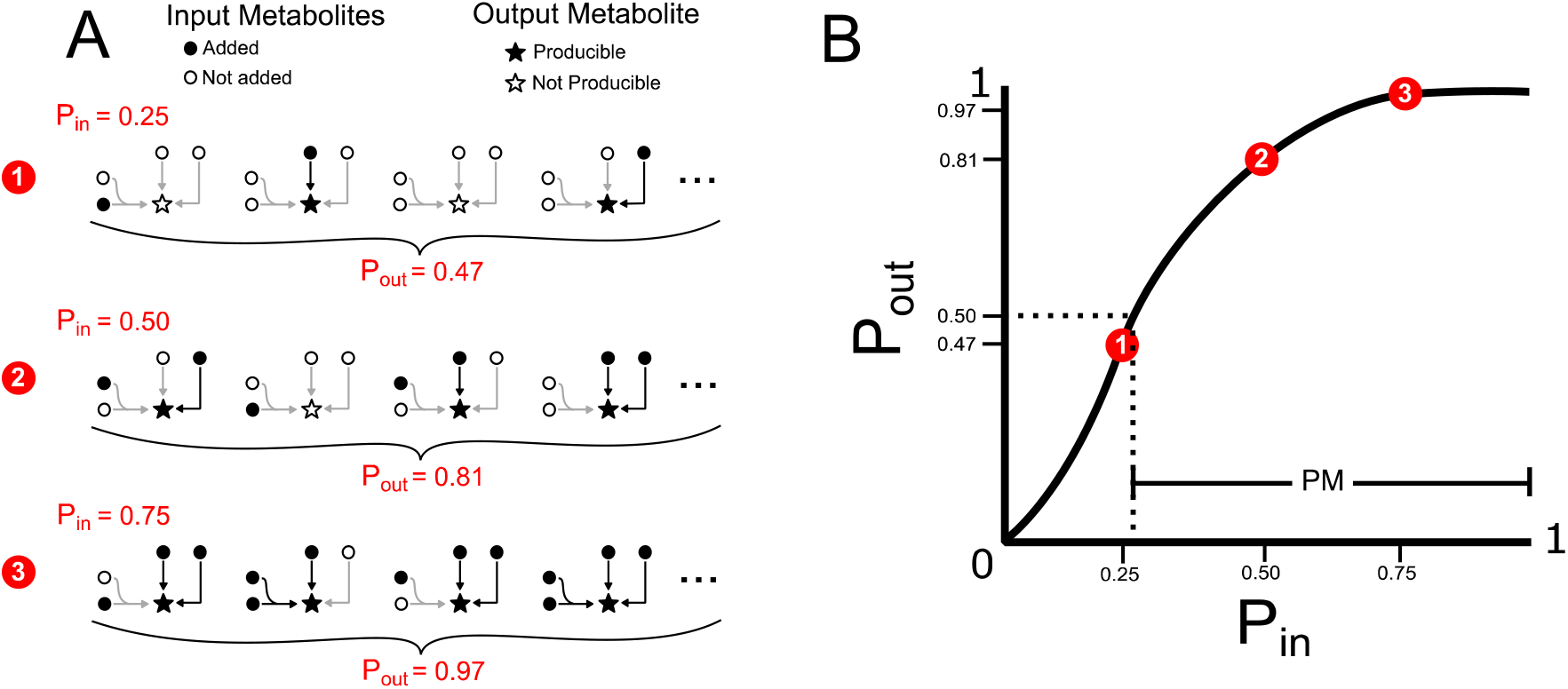
Biosynthetic network robustness analysis framework. A probabilistic framework was developed to calculate the biosynthetic network robustness of a given metabolic network and target metabolite. (A) Input probabilities (P_in_) are assigned to each input metabolite to designate the probability of adding that metabolite to the network. For our implementation, each input metabolite is assigned an identical P_in_ value. Random sets of input metabolites are sampled, based on P_in_, and a modified version of flux balance analysis is used to determine if the network can produce a specified target output metabolite for each random sample. Many random samples are taken to estimate the output probability (P_out_) of the target output metabolite. Three examples of P_m_ values and the corresponding P_out_ values are shown for a very simple network and target output metabolite. The output probabilities here were calculated using the probabilistic equation *P_out_* = 1 − [(1 − *p_in_*)^2^ * (1 − *p_in_*^2^)] = *2P_in_* – *2P_in_*^3^ + *P_in_*^4^. For more information on this equation please refer to Supplementary Figure 1. (B) A producibility curve can be calculated which represents P_out_ as a function of P_in_. Points along this curve can be sampled by assigning the P_in_ value and estimating P_out_. The three examples from A are shown in red on the curve in B. The producibility metric (PM) is used to summarize the producibility curve, and quantifies biosynthetic network robustness. It is defined by the value of P_m_ at which P_out_ equals 0.5, analogous to the K_m_ value of the Michaelis-Menten curve. PM is equal to 1 minus this value, such that increasing PM correspond to increasing biosynthetic network robustness.

The probabilistic method that we introduced allows the definition of two novel concepts, the “producibility curve” and “producibility metric” (PM) (Figure 1 B). The producibility curve is a plot of *P_out_* as a function of *P_in_*. For a given metabolic network and metabolite target, this curve can be estimated by sampling input metabolites for different values of P_in_ (between 0 and 1), and calculating P_out_ (Figure 1 B red points). The PM is a single metric which encapsulates biosynthetic network robustness by summarizing the producibility curve. The PM is defined by the P_in_ value along the producibility curve at which P_out_ is equal to 0.5. The PM value is equal to 1 minus this P_in_ value, by convention, such that larger PM values correspond to increased robustness. An analogy can be drawn between the mathematical representation of PM and the half maximal concentration constant *K_m_* in the Michaelis-Menten sigmoidal curve. Our method calculates PM efficiently by random sampling and a nonlinear fitting algorithm (for details, see methods section: Algorithm functions *calc_PM_fit_nonlin*). In addition to being quantified computationally for arbitrary metabolic networks and metabolites, the PM can also be obtained analytically by using combinatorial considerations (see Supplementary Figure 1). This analytical result clarifies the connection between our metric and the concept of minimal precursor sets^62^, and could serve as the basis for further theoretical work on the fundamental properties of metabolic networks.

The algorithms used to implement our method are written in MATLAB and designed as a set of modular functions that interface with the COBRA toolbox – a popular metabolic modeling software compendium^63,64^. The methodology behind each function is further explained in the methods section. The code is freely available online at https://github.com/segrelab/biosynthetic_network_robustness.

## Results

### Analysis of the E. coli core metabolic network

Before applying our approach to the systematic study of genome-scale metabolic networks from the human oral microbiome, we used a simpler, well characterized metabolic network model to illustrate its performance and interpretation. We applied our method to the *E. coli* core metabolic network, a simplified representation of *E. coli* metabolism consisting of central carbon metabolism and lacking peripheral metabolic pathways, such as amino acid or cofactor biosynthesis^65^. We analyzed the biosynthetic network robustness of the *E. coli* core metabolic network by calculating the PM value for all intracellular metabolites in this network. The results are shown in Figure 2 A, overlaid on the metabolic network, with each node’s color indicating its PM value and node size indicating its degree. The *E. coli* core metabolic network is highly connected and this leads to most metabolites having high PM values (PM > 0.950), matching expectations. For example, the metabolites H^+^ and pyruvate are both highly connected in the metabolic network and have high biosynthetic network robustness (PM = 0.968 and 0.952 respectively). However, the network also contains several metabolites that are well connected, but have low PM values. These include, for example, the cofactors AMP/ADP/ATP and NAD+/NADH, which have PM values of ~0.7 and ~0.5 respectively, because they can be recycled from each other, but not biosynthesized in this network. The network also includes several examples of the opposite situation, i.e. metabolites that are poorly connected but have high PM values. One example is D-lactate, which is produced via Lactate Dehydrogenase (LDH) from the high PM metabolites Pyruvate and H^+^ (Figure 2 B). This reaction also consumes NADH and produces NAD+, but because these cofactors can be easily recycled from each other by a large number of different reactions they have minimal influence on the PM value of D-lactate (Figure 2 B). This example demonstrates the fact that our metric captures metabolites which are easily produced because their precursors are easily produced, and that the utilization of recycled cofactors has minimal influence on the PM. Overall, there is no significant correlation between the PM values and the node degree of a metabolite in the network (Supplementary Figure 2), indicating that our metric describes a unique property of a metabolite in a metabolic network that is not captured simply by node degree.

**Figure 2.**
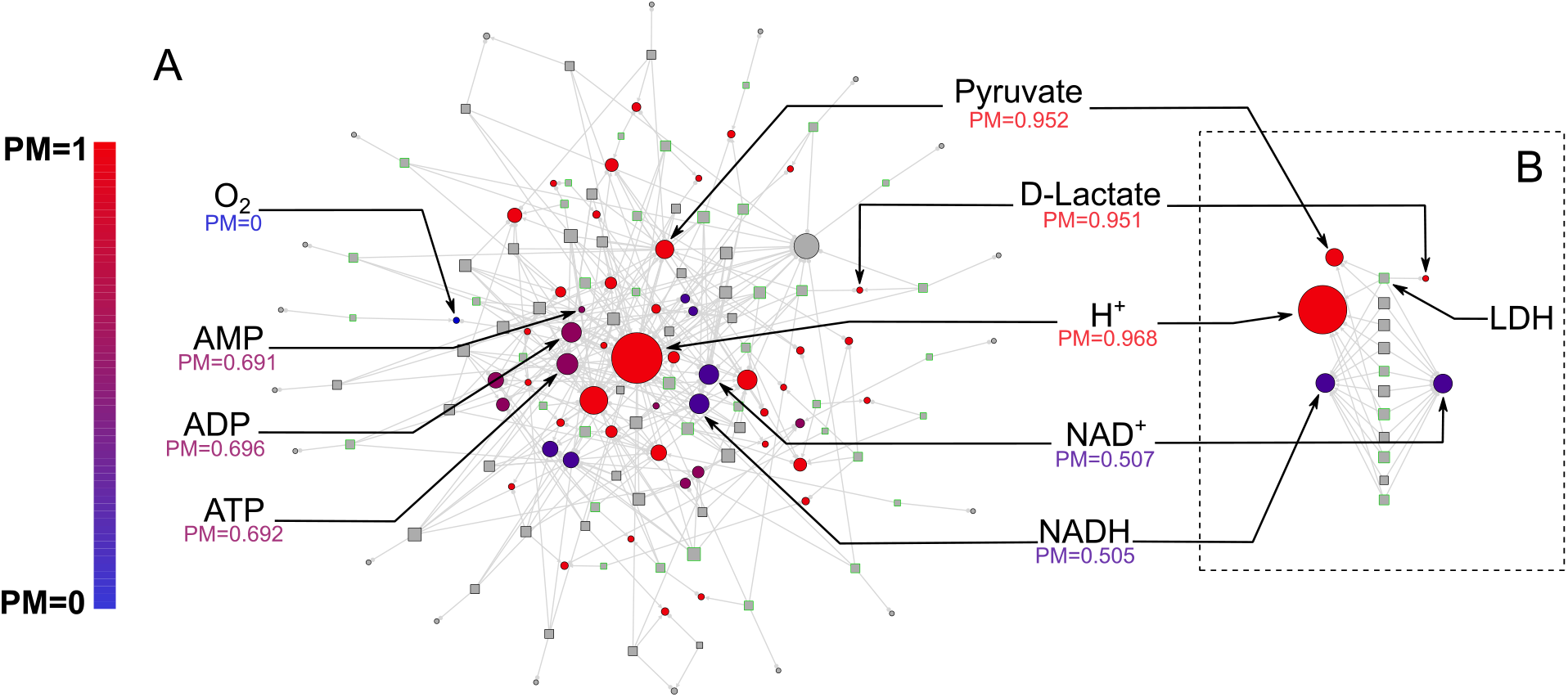
Biosynthetic network robustness of the *E. coli* core metabolic network. We calculated the producibility metric (PM) for all intracellular metabolites in the *E. coli* core metabolic network to demonstrate the implementation of our method on a simple network. (A) The network is represented as a bipartite graph with metabolites shown as circles and reactions shown as squares. Reactions shown with a green border are reversible in the model. All intracellular metabolites are colored based on their PM value (low – blue, high – red). Reactions and metabolite nodes are sized based on their total node degree. Several key metabolites of interest are highlighted with their corresponding PM values shown. Central metabolites such as H+ and Pyruvate have high degree and high PM. Cofactors such as AMP/ADP/ATP and NAD^+^/NADH have high degree but low PM, as they cannot be biosynthesized in this network. Oxygen is an example of a PM 0 metabolite that cannot be produced from any other metabolites in this network. D-lactate is an example of a metabolite with low degree and high PM i.e. it is easily produced but not well-connected. (B) Reactions related to the cofactors NAD^+^ and NADH are shown in a separate panel. The top reaction, Lactate Dehydrogenase (LDH), is shown with all substrates while all other reactions are shown without additional substrates. The metabolite D-lactate has high PM despite being poorly connected in the metabolic network because it can be produced from the high PM metabolites pyruvate and H^+^ via LDH. This reaction also consumes NADH and produces NAD^+^, however these cofactors have minimal impact on the PM because they are easily recycled in the network by a large number of different reactions.

### Reconstruction of human oral microbiome metabolic networks

We next applied our method to the human oral microbiome, aiming at a mechanistic characterization of the biochemical capabilities of different microbes based on metabolic networks reconstructed directly from their genomes. As a first step, we reconstructed metabolic networks for 456 different microbial strains representing a diverse set of human oral microbes, whose annotated genomes were available from the Human Oral Microbiome Database (see methods section for details). These organisms represent 371 different species, 124 genera, 64 families, 35 orders, 22 classes, and 12 phyla. Metadata related to the selected organisms can be found in Supplementary Table 1. Notably, the database includes several sequenced yet uncultivated or recently co-cultured organisms. This fact, together with the unique flexibility of our analysis, allowed us to obtain insight into these microbes. In particular, the following sequenced yet uncultivated, or recently co-cultured, strains were included in our analysis: *Saccharibacteria* (TM7) bacterium HMT-952 strain TM7x^25^, *Saccharibacteria* (TM7) bacterium HMT-955 strain PM004, *Saccharibacteria* (TM7) bacterium HMT-488 strain AC001, *Tannerella* HMT-286 strain W11667^66^, *Anaerolineae (Chloroflexi* phylum) bacterium HMT-439 strain Chl2^67^, *Absconditabacteria* (SR1) bacterium HMT-874 strain MGEHA^68^, and *Desulfobulbus* HMT-041 strains Dsb2 and Dsb3^69^. All of the selected genomes were used to reconstruct sequence-specific draft metabolic networks using the Department of Energy Systems Biology Knowledgebase (KBase) and the build metabolic model app^45,70,71^. The networks were reconstructed without any gap-filling to increase the specificity of the resulting predictions. A KBase narrative containing the genomes and draft metabolic network reconstructions can be found at: https://narrative.kbase.us/narrative/ws.27853.obj.935. The complete collection of all network models is also available for download in MATLAB (.mat) format at https://github.com/segrelab/biosynthetic_network_robustness.

### Large-scale analysis of biosynthetic capabilities across the human oral microbiome

We analyzed the biosynthetic network robustness for 88 different biomass metabolites across the aforementioned 456 metabolic networks from the human oral microbiome. The 88 biomass metabolites included all biomass building blocks considered to be essential for either Gram-negative or Gram-positive biomass, as listed in the KBase build metabolic models app^45,70,71^ (listed in Supplementary Table 2). Through this analysis we calculated 40,128 PM values which represent an atlas of biosynthetic capabilities across these human oral microbiome organisms. The ensuing atlas is represented as hierarchically bi-clustered PM values for all 456 organisms and 88 metabolites in Figure 3. The same data is available in Supplementary Figure 3 (clustered by taxonomy), and in Supplementary Table 3.

The hierarchically clustered heat map (Figure 3) shows extensive variability in the PM values of different organisms and metabolites across the oral microbiome. There are three main large clusters of metabolites: one cluster with consistently high PM (top), one cluster with low PM values (middle), and one cluster with variable PM (bottom). Different classes of metabolites cluster quite differently across this landscape. In addition to simple ubiquitous metabolites, such as H2O or glycine (Figure 3 I), all nucleotides have high PM across the oral microbiome organisms. Amino acids generally have high PM as well, with the notable exception of L-tryptophan (Figure 3 II). Interestingly, L-tryptophan is known to be a particularly difficult amino acid to synthesize^72^. Metal ions generally had PM value of 0 across all organisms, serving as an expected negative control. Some exceptions, such as Mg^2^+, Co^2^+, Cl^-^, Fe^3^+, and Fe^2^+, can be explained based on their presence in larger compounds, such as porphyrins. For example, Co^2^+ has increased PM values in a pattern that closely follows the PM values of the cobalt containing vitamin cobamamide (Figure 3 III).

**Figure 3.**
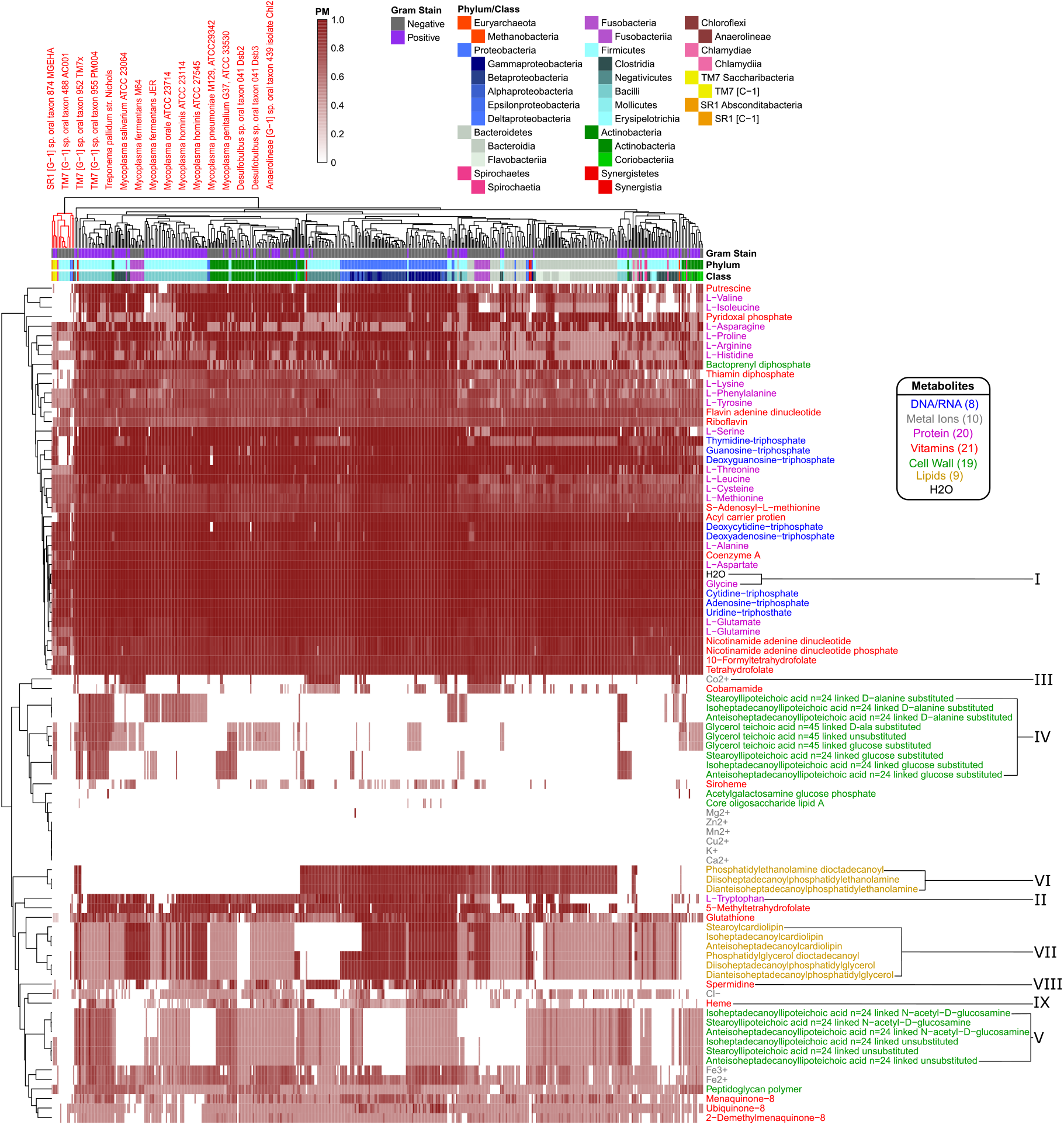
Human oral microbiome organisms biosynthetic network robustness matrix. The producibility metric (PM) was calculated for 456 different oral microbiome organisms (columns) and 88 different essential biomass metabolites (rows). The resulting matrix is hierarchically bi-clustered based on average distances between organisms and metabolites PM values. Organism Gram-stain and phylum/class are indicated by several annotation columns at the top of the matrix. The biomass metabolites analyzed consisted of several different types of metabolites indicated with different colors. Several metabolites that showed interesting patterns across oral microbiome organisms are highlighted with roman numerals. The most distinct cluster of organisms is highlighted and annotated (top left), which consisted of fastidious reduced-genome organisms *(Mycoplasma, Treponema)* and uncultivated or recently cultivated organisms *(SR1, TM7, Desulfobulbus, Anaerolineae).*

Before analyzing in detail the patterns identifiable in the PM atlas of Figure 3, we showed that such patterns cannot be trivially attributed to simple broad properties, such as genome size, even if genome size is known to be an important predictor of the overall biosynthetic capabilities of an organism^73^. Fastidious or parasitic organisms tend to have reduced genomes and consequently reduced metabolic capabilities. In our data, the overall average PM value for each organism can be partially predicted by genome size. A linear regression model and quadratic regression model which used the log of genome size to predict the average PM value across all metabolites for each organism had R^2^ values of 0.498 and 0.551 respectively (Supplementary Figure 4A). However, by using Akaike information criterion (AIC) and Bayesian information criterion (BIC) statistical analyses^74^ (Supplementary Figure. 4B, C), we found that adding taxonomic parameters to these regression models significantly improved model performance. This indicates that our data contains additional structure beyond simply genome size. In particular, both the AIC and BIC improve up to at least the order level indicating that there is additional structure up to this taxonomic level.

We further investigated, quantitatively, the associations between different taxonomic groups and the PM values of various metabolites by calculating the log likelihood ratio between a quadratic regression model predicting the PM values for a particular metabolite based solely on genome size against one that incorporates a specific taxonomic parameter of interest (Supplementary Figure 5, methods). This allowed us to highlight metabolites with highly significant increased or decreased PM values in certain taxonomic groups, and to confirm patterns that we observed by eye in Figure 3 and Supplementary Figure 3. These patterns and observations are elaborated in the following section.

### Capturing specific biosynthetic patterns across human oral microbiome organisms

Numerous patterns and details of the atlas of biosynthetic capabilities captured by the PM values (Figure 3) could be relevant for addressing specific biological questions or model refinement challenges. Here we focus in detail on two specific classes of compounds: (i) cell-wall and membrane components, which tend to vary broadly across organisms, and are important for antimicrobial susceptibility and immune system recognition; and (ii) amino acids and essential factors (e.g. vitamins), which could be relevant for understanding metabolic exchange among bacteria and with the host.

A first striking pattern in the atlas of biosynthetic capabilities captured by the PM values (Figure 3) is the complexity of cell-wall and membrane components of different taxa. Some aspects of this pattern are consistent with standard attribution of metabolites associated with the Gram staining categories (estimated using the KBase build metabolic model app^45,70,71^). However, we also observed interesting deviations, which could be partially attributed to known finer resolution in the specific membrane components across taxa. Compared to other metabolites, cell-wall components generally tend to have variable or low PM values across the oral microbiome organisms. We analyzed in detail fifteen different teichoic acids, a class of metabolites expected to be found in the cell wall of Gram-positive organisms that play an important role in microbial physiology and interactions with the host^75^. Of these, nine were found to have higher PM values in Gram-positive organisms, as expected (Figure 3 IV). In particular, the D-alanine substituted lipoteichoic acids had high PM values in the phylum *Firmicutes* and specifically the class *Bacilli.* However, there was another set of 6 teichoic acids that had intermediate PM values across a large number of organisms and didn’t follow Gram-staining trends (Figure 3 V). These consisted of three N-acetyl-D-glucosamine linked and three unsubstituted teichoic acids. As detailed in Supplementary Text 1, the increased PM for this teichoic acid in many Gram-negative species can be attributed to the presence of a specific gene^76-78^ that may merit closer inspection in the network reconstruction process.

We further observed clear trends associated with several lipids which are expected to be found in the cell membrane of both Gram-positive and Gram-negative organisms. In particular, we found a strong increase in the PM value for three phosphatidylethanolamine lipids in Gram-negative organisms (Figure 3 VI). Interestingly, these lipids have been previously observed to be more commonly produced in Gram-negative organisms, and have implications for antimicrobial susceptibility^79,80^. We also identified trends associated with three cardiolipin and three phosphatidylglycerol lipids that display generally similar PM patterns across different species (Figure 3 VII). One class of organisms that stands out with respect to lipid biosynthesis are the *Negativicutes*. These organisms have relatively high PM values for phosphatidylethanolamine but PM values of 0 for cardiolipin and phosphatidylglycerol lipids. Consistent with this result, it has been previously observed that the *Negativicutes* organism *Selenomonas ruminantium* lacks cardiolipin and phosphatidylglycerol lipids in its inner and outer cell membranes, but does have phosphatidylethanolamine^81^. It has been hypothesized that the membrane stabilizing role of these two missing lipids could be partially fulfilled by peptidoglycan bound polyamines, including spermidine, in *Selenomonadales* organisms^81,82^. Concordantly, we see an increased PM value for the polyamine spermidine across *Negativicutes* in our data (Figure 3 VIII).

Aside from lipids and cell-wall components, there are a number of interesting trends related to several amino acids and other essential factors in our data. A number of metabolites had notably increased PM in the phylum *Proteobacteria* and decreased PM values in the phylum *Bacteroidetes*. A notable example is heme, which can be seen to follow this trend (Figure 3 IX). Heme plays an important role in microbe host interactions, as bacterial pathogens often acquire it from their human host^83^. In the context of the human oral microbiome, the oral pathogen *Porphyropmonas gingivalis* (belonging to the class *Bacteroidetes*) is known to scavenge heme^84^, compatible with the above pattern. Other metabolites that displayed the same trend include: L-arginine, L-cysteine, L-methionine, L-tryptophan, and glutathione. L-arginine can be catabolized via the arginine deiminase pathway to regenerate ATP and is thus an interesting exchange metabolite beyond its use as a protein building block^85,86^. L-tryptophan is one of the highest cost amino acids to biosynthesize^72^, and thus is an intriguing exchange candidate. L-methionine and L-cysteine are the only two sulfur containing standard amino acids, and glutathione is synthesized from L-cysteine. It’s possible that the discrepancies between PM values observed here are indicative of broad amino acid and vitamin exchange between the classes *Proteobacteria* and *Bacteroidetes* in the human oral microbiome.

### Uncovering biosynthetic deficiencies in fastidious human oral microbiome organisms

In addition to dissecting the patterns associated with specific metabolites, one can analyze the PM landscape of Figure 3 from the perspective of the organisms and their agglomeration into clusters. Given their importance in disease and the unresolved challenges related to their reduced metabolic capabilities, we focused specifically on fastidious human oral microbiome organisms. Strikingly, in our large clustered PM matrix, the most distinct hierarchical cluster of organisms consisted of a number of fastidious organisms (Figure 3 top left). This cluster included all of the *Mycoplasma* genomes that we analyzed, and one *Treponema* genome. *Mycoplasma* and *Treponema* are genera that are known to be parasitic and have evolved to have reduced genomes and metabolic capabilities^87-91^. The remaining members of this cluster included nearly all of the sequenced yet uncultivated, or recently co-cultured, organisms in our study. The organisms included were from the phyla: *Absconditabacteria* (SR1), *Saccharibacteria* (TM7), *Proteobacteria* (genus *Desulfobulbus),* and *Chloroflexi* (class *Anaerolineae).* Only one of the previously uncultivated organism we analyzed was found outside of this fastidious cluster, namely *Tannerella* HMT-286. Interestingly, this bacterium is hypothesized to rely on externally supplied siderophores to support its growth^66^. This type of protein dependency is not captured by our metabolic analysis and highlights the fact that, while uncultivability can be driven by many different mechanisms, our method captures the prominent effect of reduced metabolic capacity. The other uncultivated organisms that we identified in this cluster have been hypothesized to have reduced genomes and limited metabolic capabilities underlying their fastidious nature, much like *Mycoplasma*.

We sought to gain clearer insight into the metabolic properties of these co-clustered fastidious organisms by re-clustering their PM submatrix (Figure 4 A). By comparing the PM values in this fastidious cluster to those in the average oral microbiome organisms, it is clear that the fastidious organisms had reduced PM values for a large number of metabolites including cell-wall components, lipids, amino acids, and other essential factors. When ranking metabolites by their difference in average PM between all oral microbiome organisms and the fastidious cluster a number of amino acids and vitamins stand out as being the most depleted in the fastidious cluster. The top metabolites where: pyridoxal phosphate, L-valine, putrescine, L-isoleucine, bactoprenyl diphosphate, thiamin diphosphate, 5-methyltetrahydrofolate, L-lysine, deoxyguanosine triphosphate, L-tryptophan, and guanosine-triphosphate. These metabolites may be particularly relevant with regards to exchange between fastidious organisms and their oral microbiome community partners. Amino acids, in particular, have been hypothesized to be involved in metabolic exchange between microbial organisms in communities^1, 7 37,92^ Notably, amino acids with reduced PM in the fastidious cluster (i.e. amino acids more readily produced by other organisms) tend to be among the more costly ones^72^, as indicated by a Spearman correlation analysis (ρ = 0.4595, P-value = 0.0415). An exception to this trend, potentially interesting for follow up studies, is the case of the branched chain amino acids L-valine, and L-isoleucine, which are the two amino acids with most reduced PM in fastidious organisms, but are not among the costliest. Notably, branched chain amino acid supplementation has been shown to alter the metabolic structure of the gut microbiome of mice^93^.

**Figure 4.**
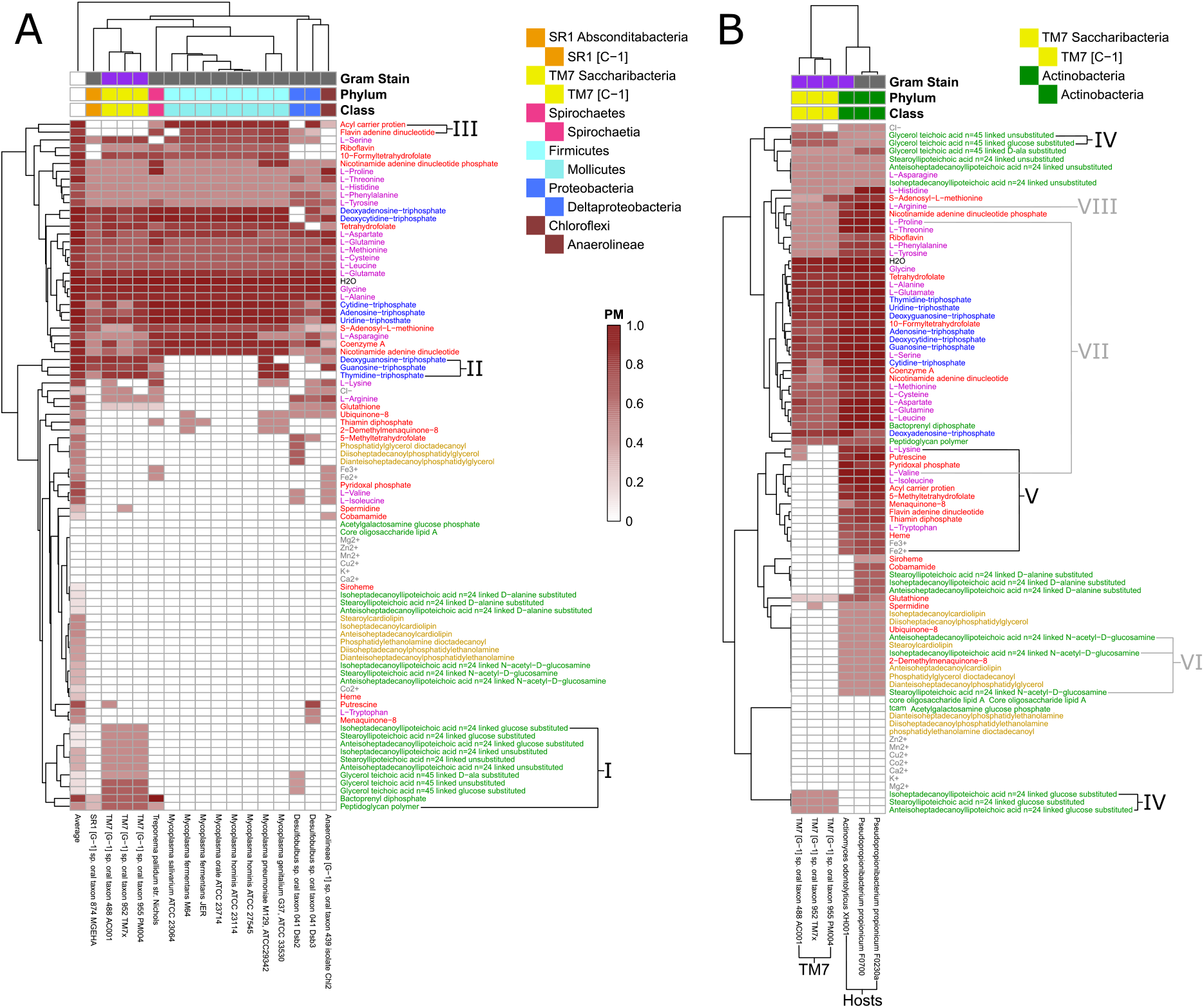
Biosynthetic network robustness sub-matrices for fastidious/uncultivated and TM7/host organisms. Sub-matrices of the larger biosynthetic network robustness matrix were re-clustered to highlight variations within specific groups of fastidious and uncultivated organisms. (A) The fastidious/uncultivated organisms that were identified as the most unique cluster in the larger matrix from Figure 3 were re-clustered hierarchically based on average distance between organisms and metabolites producibility metric (PM). The average PM value across all oral microbiome organisms analyzed in this study is shown in the far left column. Differences between the fastidious *Mycoplasma* genus and the previously uncultivated TM7 species are highlighted. (B) The PM values for the previously uncultivated TM7 species and their co-culture growth-supporting hosts bacteria were extracted and re-clustered hierarchically based on average distance between organisms and metabolites PM values. Differences between the TM7 species and their bacterial hosts are highlighted.

To gain more specific insight into a specific class of recently-cultivated fastidious organisms, *Saccharibacteria* (TM7), we further focused our analysis on identifying discrepancies between *Mycoplasma* and TM7. Our analysis included eight *Mycoplasma* genomes and three TM7 genomes. *Mycoplasma* are a relatively well characterized genus of intracellular parasites with reduced metabolic capabilities, and TM7 are a recently co-cultured phylum of the candidate phyla radiation that display reduced metabolic capabilities and a parasitic lifestyle. Comparing these two groups of organisms gives deeper insight into the unique metabolic capabilities of each. There are several cell-wall components for which TM7 has relatively high PM values and Mycoplasma has PM values of zero (Figure 4 I). These include nine different teichoic acids, bactoprenyl diphosphate, and peptidoglycan. This highlights extensive cell-wall/peptidoglycan metabolism in TM7 organisms and the known lack of a cell-wall in *Mycoplasma^91^.* Furthermore, a set of three nucleotides: dGTP, GTP, and TTP, have high PM values for TM7 and PM values of zero for *Mycoplasma* organisms (Figure 4 II). This pattern of nucleotide biosynthesis deficiency in *Mycoplasma* is consistent with the observation that some strains have been shown to be dependent on supplementation of thymidine and guanosine but not adenine or cytosine nucleobases for growth^94^. Finally, the cofactors acyl carrier protein (ACP) and flavin adenine dinucleotide (FAD) had high PM values in *Mycoplasma* and PM values of zero in TM7 organisms (Figure 4 III). The lack of these cofactors in TM7 seems surprising, but is indeed matched by a complete lack of any metabolic reactions annotated to utilize FAD and ACP as cofactors in the draft reconstruction of the TM7 metabolic networks.

In addition to investigating the metabolic deficiencies of fastidious organisms, the PM landscape gave us the opportunity to compare these gaps with possible complementary capabilities in organisms known to support their growth. The three TM7 strains that we analyzed were recently co-cultured with host bacteria from the human oral microbiome. TM7x was shown to be a parasitic epibiont of *Actinomyces odontolyticus* XH001 ^25,26,95^. TM7 AC001 and PM004 were recently both co-cultured successfully with either of the host strains *Pseudopropionibacterium propionicum* F0230a or F0700 (not yet published). We sought to further investigate these newly discovered relationships to gain insight into possible metabolic exchange (Figure 4 B). Interestingly, TM7 organisms had higher PM values than their host strains for several cell-wall components: three glucose-substituted teichoic acids, and glucose-substituted and unsubstituted glycerol teichoic acid (Figure 4 IV), suggesting that TM7 is capable of producing several cell-wall components that its host cannot. Conversely, as expected, a large number of metabolites had increased PM values in the host strains compared to the TM7 strains. These metabolites are hypothesized to be easily synthesized by the host and not TM7 and are thus interesting candidates for growth supporting exchange in co-culture. Fourteen different metabolites had average PM values in the hosts greater than 0.60 higher than in the TM7 organisms (Figure 4 V). The ranked list includes: L-isoleucine, L-valine, acyl carrier protein, 5-methyltetrahydrofolate, pyridoxal phosphate, flavin adenine dinucleotide, thiamin diphopsphate, putrescine, L-tryptophan, Fe^2^+, heme, Fe^3^+, L-lysine, and menaquinone-8. Interestingly, the branched chain amino acids L-isoleucine and L-valine are again at the top of the list. The correlation of amino acid biosynthesis cost^72^ with the difference in PM values between host and TM7 is even higher than what we observed across all fastidious organisms (Spearman correlation ρ = 0.6011, P-value = 0.0051).

Our results provide context and putative mechanistic details related to observed gene expression and metabolic changes in TM7-host co-culture. In particular, the first and currently only published work on co-culture involving TM7 is the one on TM7x with the host *Actinomyces odontolyticus* XH001 ^25,26,95^. Transcriptomic data for the co-culture of TM7x and *A. odontolyticus* XH001 showed that a number of genes associated with N-acetyl-D-glucosamine were up regulated in *A. odontolyticus* in this interaction. Our results show that, although TM7 does have extensive cell wall metabolism, *A. odontolyticus* has higher PM for N-acetyl-D-glucosamine substituted components (Figure 4 VI). This suggests that the host is responsible for the biosynthesis of these cell-wall components, which may be overexpressed in co-culture. Metabolomics experiments from this co-culture have identified the cyclic peptide cyclo(L-Pro-L-Val) as a potential signaling molecule in this relationship. Our PM analysis suggests that this molecule would be synthesized by the host as it has increased PM values for both of the amino acids included (Figure 4 VII). In fact, L-valine has one of the highest discrepancies in PM for host and TM7. Finally, another potentially exchanged amino acid of interest is L-arginine. All three TM7 draft metabolic network reconstructions that we analyzed were annotated to possess either all or all but one of the reactions in the arginine deiminase pathway (TM7 PM004 is missing the arginine iminohydrolase reaction) (See also supplementary figure 6 and interactive Cytoscape^96^ file for a representation of the full metabolic network for each TM7 strain including PM calculations for all intracellular metabolites and subnetworks of the arginine deiminase pathway, Supplementary Files 1-3). This catabolic pathway can be used to degrade L-arginine to regenerate ATP, and has been implicated in syntrophic microbial interactions^85,86^. In our PM analysis L-arginine had consistently higher PM in host than TM7 (Figure 4 VIII). Thus, L-arginine exchange and metabolism via the arginine deiminase pathway could contribute to the dependence of TM7 on its hosts (Figure 5).

**Figure 5.**
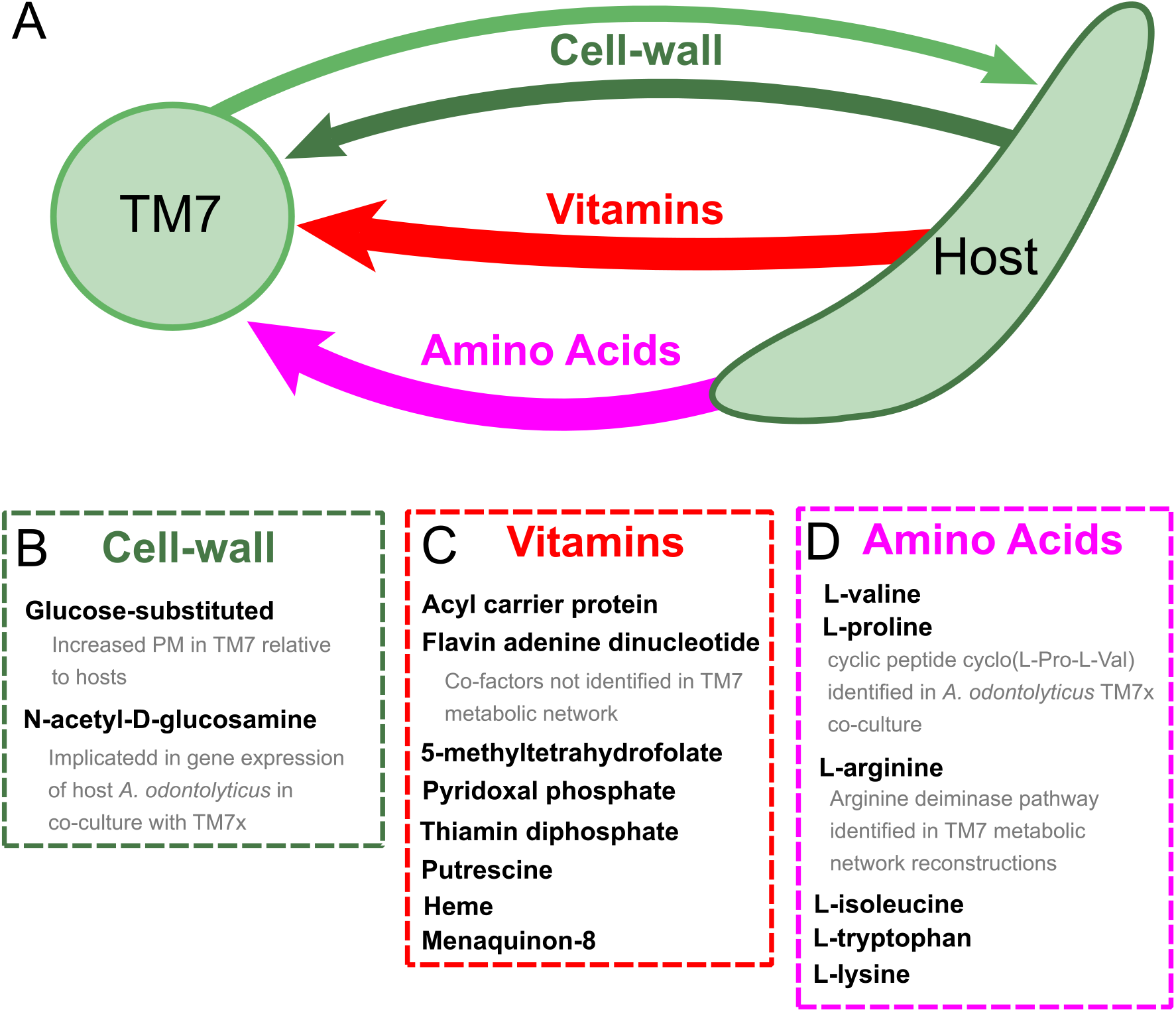
Hypothesized metabolic exchange between TM7 and their bacterial hosts. (A) Hypotheses were generated regarding the exchange of metabolites between TM7 species and their bacterial hosts by comparing their producibility metric (PM) across essential biomass metabolites. Many metabolites of different types were observed to have higher PM values in one set of organisms when compared to the other (arrows point from high to low). (B) The cell-wall components containing glucose-substituted teichoic acids were among the only metabolites with PM higher in TM7 than in hosts. N-acetyl-D-glucosamine-substituted teichoic acids had increased PM in the host relative to TM7, and previous gene expression data from the co-culture of TM7x and *A. odontolyticus* shows several genes related to N-acetyl-D-glucosamine that are overexpressed in *A. odontolyticus* during co-culture^25^. (C) Several vitamins/cofactors/other essential factors had significantly decreased PM in TM7 compared to the hosts. The cofactors acyl carrier protein and flavin adenine dinucleotide had decreased PM in TM7, and were also not found to be utilized in the TM7 draft metabolic network reconstructions. (D) Several amino acids had significantly decreased PM in TM7 compared to the hosts. L-valine and L-proline were both decreased in TM7 relative to the host, and previous metabolomics data from the co-culture of TM7x and *A. odontolyticus* identified the cyclic dipeptide cyclo(L-Pro-L-Val) as a potential signaling molecule^25^. L-arginine had decreased PM in TM7 relative to the host and could potentially be exchanged and catabolized by TM7 via the arginine deiminase pathway.

## Discussion

We have developed a novel method for analyzing the biosynthetic capabilities of microbial organisms based on draft metabolic networks reconstructed directly from genomic information. Our method provides a preliminary assessment of the biosynthetic capabilities of a metabolic network model, without the need for gap-filling, that can be used to gain biological insight and evaluate initial model performance. The concept we define, biosynthetic network robustness, provides an environment-independent evaluation and utilizes all available stoichiometric constraints. Environmental independence is achieved by randomly sampling many possible nutrient combinations in a probabilistic manner and computing a metric inspired by percolation theory. This measure defines the robustness with which an organism can produce a given metabolite from any random set of precursors and thus avoids the issue of metabolite producibility being inherently dependent on environment^49,50^. In this work we have chosen to calculate the metabolic properties of organisms without assuming a particular environment; however, future implementations could utilize environmental information in a probabilistic manner when appropriate. Additionally, we have analyzed the production of individual target metabolites, but our method could easily be extended to sets of metabolites such as the simultaneous production of all biomass components. Our method utilizes all available stoichiometric constraints of the metabolic network as opposed to an adjacency matrix used by alternative approaches^48^. Stoichiometric constraints are implemented with a modified version of flux balance analysis (See methods section algorithm functions: *feas*), as opposed to the alternative network expansion algorithm to avoid the dependence on cofactors as bootstrapping metabolites^97^.

It is important to highlight that several assumptions are made in the representation of enzymatic reactions as a network that generally limit metabolic network analysis methods. The primary limitation is in enzyme annotation. Aside from missing or incorrect annotations, subtle processes such as enzyme promiscuity and spontaneous reactions may have unquantified effects on metabolic network function. Reaction direction/reversibility is also difficult to predict as it requires detailed knowledge of reaction thermodynamics and metabolite concentrations. In particular, inaccurate or missing information about reaction direction/reversibility could lead to uncertainty about whether a high PM from our method should be interpreted as reflecting biosynthetic or degradative capabilities (or both). Throughout our analysis we have utilized default reversibility constraints provided by the KBase build metabolic models app^45,70,71^, but more stringent constraints on directionality could possibly improve our results. Additionally, as our method analyzes local properties of the metabolic network (the PM value for a specific metabolite) unidentified gaps in biosynthetic pathways that occur in close proximity to the target metabolite of interest could lead to incorrect predictions regarding microbial auxotrophy. In general, all metabolic network analysis methods face similar limitations. Even as newly developed experimental methods gradually improve metabolic reaction annotation^98-101^, it is likely that we will have to continue dealing with incomplete knowledge. Thus, approaches such as ours are valuable for initial assessment of metabolic capabilities with minimal arbitrary assumptions, and unexpected modeling results can help to pinpoint specific areas in need of refinement.

In applying our method to the human oral microbiome, we computed an atlas of biosynthetic capabilities across organisms that can be mined for relevant biological insight. Overall, many of our predictions were consistent with known patterns such as the reduction in biosynthetic capabilities in the genus *Mycoplasma* or the distribution of lipids and cell-wall components in Gram-positive and negative organisms. Additionally, unexpected predictions served as opportunities to highlight novel biological patterns or emphasize areas of the metabolic network that merit additional attention in the network reconstruction process. Our focus was on fastidious and uncultivated organisms in particular, and using our method we highlighted a unique cluster of such organisms with reduced biosynthetic capabilities. This cluster included three previously uncultivated *Saccharibacteria* (TM7) phylum organisms that were recently successfully cocultured with growth supporting bacterial host organisms. Our method singled out specific biosynthetic capabilities of these organisms, and was used to develop hypotheses regarding metabolic exchange between TM7 and host bacteria that give context to existing co-culture data and should be further testable in future experiments. These three TM7 species are the first successfully cultured organisms from the candidate phyla radiation and therefore are of general interest beyond their role in human oral health. In fact, the recent identification of the candidate phyla radiation demonstrates the broad prevalence across the tree of life of reduced-genome organisms that potentially rely on their community context for metabolic supplementation^15-17^. Further analysis of these organisms with our method could continue to provide insight into their unique metabolic properties.

By quickly translating genotype into phenotype with minimal assumptions, our approach has the potential to serve as a baseline estimate of metabolic mechanisms in different microbial communities and allows us to more easily decipher microbial community structure and function. Our method can be easily applied other human-associated or environmentally relevant microbial communities, providing valuable putative insight into inter-microbial metabolic dependencies, that could be used to interpret existing data or design future experiments. In particular, we envisage that this type of metabolic insight could help bridge the gap between correlation studies and a mechanistic understanding of microbial community metabolism and dynamics.

## Methods

### Method implementation

The framework for implementing our method was developed as several different modular functions that interact in a nested manner to run our analysis. The structure of these functions and their associated variables is described in Supplementary Figure 7 via a code diagram. The functions are written in MATLAB and interface with the COBRA toolbox^63,64^. The code is built around the COBRA toolbox commands *changeObjective* and *optimizeCbModel*. Thus, running our code requires installation of the COBRA toolbox. Additionally, the nonlinear fitting function utilizes the MATLAB function *lsqnonlin* for nonlinear least squared fitting. Additional functions were developed to implement our probabilistic framework and run our analysis method. Any of these functions could be replaced with alternative modules that improve or expand upon the algorithm in the future. We describe here each modular function, providing details on the computations performed. The full code for implementing our method is available online at https://github.com/segrelab/biosynthetic_network_robustness.

#### Algorithm functions

##### feas

This function determines if the production of a given target metabolite set is feasible given the metabolic network model with specified constraints. Flux balance analysis was used to determine the feasibility of production^46^. Flux balance analysis was chosen over the alternative network expansion algorithm due to its treatment of cofactor metabolites^97^. In network expansion, cofactors must be added to the network to “bootstrap” metabolism, whereas in flux balance analysis any reaction utilizing a cofactor can proceed given that the cofactor can be recycled by a different reaction, which is a less restrictive constraint on the metabolic network flux. Furthermore, our implementation allows for inequality or equality mass balance constraints. Traditional flux balance imposes an equality mass balance which is often referred to as a steady state constraint. This constraint restricts the rate of change of all metabolite concentrations to be equal to 0. We provide the option of implementing inequality mass balance, which constrains the rate of change of metabolite concentrations to be greater than or equal to 0. In practice, inequality mass balance is implemented by adding unbounded exporting exchange reactions and calculating steady state solutions. We have implemented inequality mass balance for all of our calculations due to the fact that we are analyzing local properties of the metabolic network (the production of a single metabolite) and do not want the network to be constrained by the global requirement to achieve steady state. During the production of a particular metabolite, the metabolic network is thus free to produce byproducts that are used elsewhere or secreted. To determine production feasibility, the export of a particular target metabolite is set to the objective function and maximized. If the maximal flux is greater than a hard-coded threshold (>0.001), then the target metabolite is considered to be feasibly produced. This function uses the COBRA commands *changeObjective* and *optimizeCbModel* to set and maximize the appropriate objective function. Mathematically, flux balance analysis is implemented as a linear programming problem with the following definition:

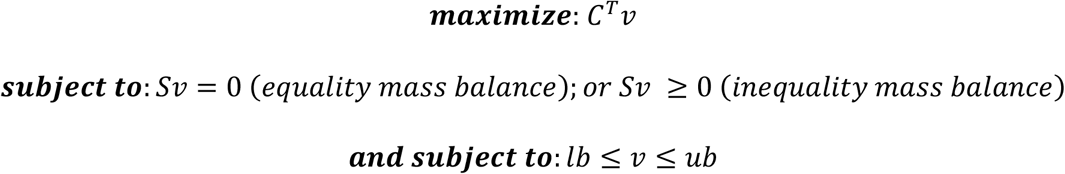

Where: *C^T^* is the transpose of a column vector indicating which reactions are to be maximized.

In this case, this specifies the exporting exchange reactions corresponding to the target metabolites. *v* is a column vector of metabolic reaction fluxes. *S* is the stoichiometric matrix describing the reactions present in the metabolic network. This is a metabolites by reactions size matrix. Each element in the matrix is the stoichiometry of a particular metabolite associated with a particular reaction. Negative values indicate that a metabolite is a reactant of that reaction being consumed, while positive values indicate that a metabolite is a product of that reaction being produced. *lb* and *ub* are the lower and upper bounds of all reactions, which define reaction reversibility or are set to −1000 and 1000 respectively when unbounded. Additional information on flux balance analysis can be found in this publication describing its implementation in detail^46^.

##### rand_add

This function is designed to give a random sample of input metabolites to be added based on the Bernoulli parameter for each input metabolite. This function uses the MATLAB *rand* function to choose a random number between 0 and 1 for each input metabolite. If this number is less than the Bernoulli parameter for that input metabolite, then the metabolite is added.

##### prob

This function utilizes *rand_add* and *feas* to determine the probability of producing the target metabolite given the input metabolite Bernoulli parameters, the metabolic network structure, and the specified constraints. A chosen number of random samples of input metabolites are generated by repeatedly running the *rand_add* function. The probability of producing the target metabolite is determined as the number of feasible trials divided by the total number of samples. The default number of samples used for the bulk of the analysis in this work was 50.

##### calc_PM_fit_nonlin

This function calculates the PM for a specified metabolic network model and metabolite using an efficient nonlinear fitting technique. The nonlinear fitting algorithm estimates the PM by randomly sampling points on the PC that fall near PM. The algorithm starts by sampling a point in the middle of the PC and then using the MATLAB function *lsqnonlin* to fit a sigmoidal curve to the sampled points of the PC. The fit sigmoidal curve is used to estimate the PM. Next, a new sample point is obtained which is offset from the estimated PM value with some noise introduced with the specified noise parameter. In this way the algorithm converges on the PM value and samples points around PM, thus increasing the accuracy of its estimate with each iteration. The estimate converges when a specified n estimates of the PM value are all within a specified threshold. The code allows for a figure to be displayed which shows the sampled data points and fit sigmoidal functions, which is useful for debugging the algorithm and finding suitable parameters. The default parameters, associated with this function, used for the bulk of our analysis were: noise = 0.3, n = 7, thresh = 0.01. The parameters chosen were selected by hand to provide good performance.

##### prep_mod

This function is used to prepare the metabolic network model for analysis with our method. The input for this function is a COBRA model, which is saved as a MATLAB structure variable. This code has been developed and optimized to work with KBase generated metabolic networks and is not guaranteed to work with networks from other sources that have different naming conventions. The first modification to the networks is to find and turn off all exchange and maintenance reactions to standardize the network models. Second, the extracellular and intracellular metabolites are identified based on naming conventions and output from the function. Third, exchange reactions are added for each metabolite (producing 1 unit of that metabolite), and a vector indicating the mapping from metabolites to these exchange reactions is output from the function. This vector is used by our method to control the presence and absence of input metabolites in the network model as well as to adjust the inequality mass balance constraints. The final output is a new network model which has been standardized for our method and in which the presence and absence of metabolites can be easily manipulated.

##### find_PM_mods_mets

This function is designed to facilitate the parallelization of the PM calculation. The function takes as inputs a directory of metabolic network models, a directory to store results, a list of target metabolite names, the index of the current network model and metabolite being analyzed and all of the specifications necessary for running *calc_PM_fit_nonlin*. The metabolite and model being analyzed can be changed dynamically to allow for parallelization. In addition to these inputs, this function has several inputs that allow for standard modifications to the PM calculation procedure. It allows for certain metabolites to be fixed on or off. It allows for several choices of metabolites to be added during the PM calculation process, including adding all intracellular or extracellular metabolites and including the target metabolite or not. It also allows for specification of the inequality mass balance constraint as either all metabolites set to inequality mass balance or all metabolites set to equality mass balance. Furthermore, it has a parameter for the number of runs to calculate the PM to obtain statistics regarding the variability of *calc_PMfit_nonlin.* For the analysis done in this work: calculation of PM for single metabolites was done by adding all intracellular metabolites (excluding targets). The mass balance constraint was set to use inequality constraints for all metabolites. The number of runs was set to 10.

#### Parallelization

We used the Boston University shared computing cluster to run our analysis for a large number of metabolic networks and metabolites. The calculation of the PM for each individual network model and metabolite can be run in parallel, vastly increasing the number of possible computations. The average runtime for computing the PM for an individual network and metabolite for 10 repeated runs was ~9 minutes and the maximum run time was ~45 minutes, given the default parameters used in this study: a = 0, s = 1, samp = 50, noise = 0.3, n = 7, thresh = 0.01, runs = 10.

#### Analysis of the E. coli core metabolic network

Our analysis method was initially demonstrated on the *E. coli* core metabolic network. We used the network provided by the BiGG database^102^. We calculated the PM value for each intracellular metabolite. The input metabolites for our PM calculations were assigned as all intracellular metabolites. This was the most naïve assumption we could use for assigning input metabolites. Additionally, using intracellular metabolites as input metabolites avoids errors that could arise from poorly annotated transporters in draft metabolic network reconstructions. Calculations were performed using the Boston University shared computing cluster to parallelize runs across networks and metabolites and improve computation time. The results of our simulation were visualized using the Cytoscape network visualization software^96^. The entire *E. coli* core metabolic network is shown, excluding the biomass reaction for clarity.

#### Reconstruction of human oral microbiome metabolic networks

A set of 456 draft metabolic networks were reconstructed for oral microbiome strains. Strains were chosen to match the sequences chosen for dynamic annotation on HOMD which cover at least one strain for each sequenced species and repeated strains for sequences of particular interest for the human oral microbiome. Several strains were additionally selected due to our interest in fastidious and uncultivated organisms. These included 8 uncultivated or recently cocultured strains. When considering the taxa TM7 and *Tannerella* sp. oral taxon 286, we chose to include the most recent genome sequences from oral microbiome co-culture experiments, although there are several additional single-cell and metagenome assembled sequences also available for *Tannerella* sp. oral taxon 286 and TM7 in particular^15, 29,103-105^. The host strains *Actinomyces odontolyticus* XH001, *Pseudopropionibacterium propionicum* F0700, and *Pseudopropionibacterium propionicum* F0230a were included due to their support of TM7 organisms in co-culture. All genomes were either found in the KBase central data repository or manually annotated with RAST and uploaded to KBase^70,71,106,107^. Strains that were dynamically annotated on HOMD but could not be found on KBase, were not of interest due to uncultivability, and already had a representative strain from their matching species were not included in our set of strains. Several naming discrepancies existed between KBase and HOMD, which are highlighted in the KBase download notes column of Supplementary Table 1. All metabolic networks were reconstructed using a KBase narrative containing all of the genomes and metabolic networks from this work, which is available to be copied, viewed, edited, or shared at https://narrative.kbase.us/narrative/ws.27853.obj.935. Metabolic networks were reconstructed for each strain with automatic assignment of Gram-stain, and without gap-filling. Metabolic network reconstructions were then downloaded from KBase as SBML files and converted to COBRA .mat files using the COBRA command *readCbModel.* Metadata related to all organisms and metabolic networks are available in Supplementary Table 1.

#### Large-scale analysis of biosynthetic capabilities across the human oral microbiome

We investigated the large-scale biosynthetic properties of the human oral microbiome by analyzing reconstructed metabolic networks for 456 different oral microbiome strains. For each metabolic network we calculated the PM value for 88 individual biomass components (40,128 total PM calculations). The biomass components were chosen to be the union of the set of default KBase Gram-positive and Gram-negative biomass compositions (see Supplementary Table 2 for details). The metabolites sulfate and phosphate were not included, while the metabolite H2O was included as a positive control. The calculations were parallelized across metabolic networks and metabolites using the Boston University shared computing cluster to improve computation time. The PM values were stored as a matrix of organisms by metabolites PM values. This matrix was analyzed using hierarchical bi-clustering based on average differences between groups. The matrix was clustered and visualized using the R package *pheatmap*.

For the comparison of average PM values and genome size, genome size was taken from KBase and added to Supplementary Table 1. We used regression modeling to identify the broad relationship between genome size, taxonomy, and the average PM value. We fit PM values to linear and quadratic models of log genome size:

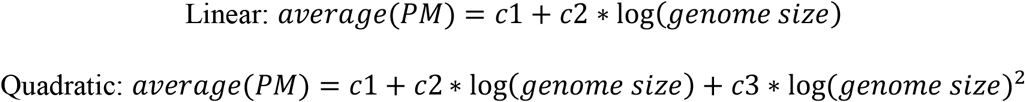

Nominal taxonomic parameters were added to these models to determine if they could improve the models prediction of PM values. Gram-stain was assigned based on KBase default assignments. Phylum, and Class were assigned based on human oral microbiome database taxonomy annotations. Regression models were developed using the MATLAB command *fitlm.* The AIC and BIC were calculated to assess model improvement upon subsequent addition of taxonomic parameters by determining if the likelihood of the model was improved while including a penalty term for each additional independent variable. Independent variables were added for each additional nominal parameter added (for example: adding the predictor of phyla meant adding 12 independent variables, one for each different phylum). The AIC and BIC were calculated using the MATLAB command *aicbic*.

#### Capturing specific biosynthetic patterns across human oral microbiome organisms

We investigated specific trends in metabolite PM values related to taxonomy by analyzing the clustered matrix of PM values. Additionally, a similar regression model was used to provide quantitative insight. The base model was a quadratic model using the log of genome size as the predictor of the specific PM value for a certain metabolite across all organisms:

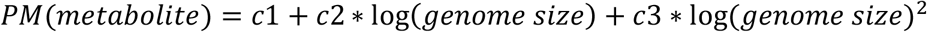

Nominal taxonomic parameters were added one at a time. Taxonomic parameters of Gram-stain (+ or -), phylum (belonging to 1 of 12 phyla or not) and class (belonging to 1 of 22 classes or not) were used. We calculated the log likelihood ratio by taking difference between the log likelihood of the base quadratic model of genome size and the model including a specific taxonomic parameter. We identified highly significant relationships using an alpha value of 10^-6^ and Bonferroni correction for multiple hypothesis testing.

#### Uncovering biosynthetic deficiencies in fastidious human oral microbiome organisms

A subset of fastidious organisms identified from the larger clustered matrix of all oral microbiome organisms PM values were re-clustered and analyzed further. The clustering method used was the same as for the larger Figure 3. Additionally, three previously uncultivated TM7 organisms (TM7x, AC001, and PM004) and several host strains for the uncultivated TM7 *(Actinomyces odontolyticus* XH001, *Pseudopropionibacterium propionicum* F0700, and *Pseudopropionibacterium propionicum* F0230a) were re-clustered and analyzed. Metabolites were ranked and analyzed based on the difference between the average PM value of separate groups. Three different ranking were used throughout this analysis 1) average fastidious cluster organisms PM subtracted from average oral microbiome organisms PM 2) average *Mycoplasma* PM subtracted from average TM7 PM 3) average TM7 host PM subtracted from TM7 PM. Correlations between amino acid biosynthetic cost^72^ and difference in PM were calculated using Spearman’s rank correlation and the MATLAB command *corr.*

## Acknowledgments

We would like to acknowledge our collaborators at the Forsyth institute for providing valuable knowledge and insight into the human oral microbiome and uncultivated microorganisms. We would also like to acknowledge all of the members of the Segrè lab and Daniel Sher of Haifa University for helpful discussions and comments on this work. Research reported in this publication was supported by The National Institute of Dental and Craniofacial Research of the National Institutes of Health under award numbers R37DE016937, R01DE024468, by National Institutes of Health grants R01GM121950 and Sub_P30DK036836_P&F, by the Defense Advanced Research Projects Agency (Purchase Request No. HR0011515303, Contract No. HR0011-15-C-0091), the U.S. Department of Energy (DE-SC0012627), the Boston University Interdisciplinary Biomedical Research Office, and by the Boston University training program in quantitative biology and physiology under Ruth L. Kirschstein National Research Service Award T32GM008764 from the National Institute of General Medical Sciences. The content is solely the responsibility of the authors and does not necessarily represent the official views of the granting agencies.

